# Joint antibiotic and phage therapy: addressing the limitations of a seemingly ideal phage for treating *Staphylococcus aureus* infections

**DOI:** 10.1101/2020.04.24.060335

**Authors:** Brandon A. Berryhill, Douglas L. Huseby, Ingrid C. McCall, Diarmaid Hughes, Bruce R. Levin

## Abstract

In response to increasing frequencies of antibiotic-resistant pathogens, there has been a resurrection of interest in the use of bacteriophage to treat bacterial infections: phage therapy. Here we explore the potential of a seemingly ideal phage, PYO^Sa^, for combination phage and antibiotic treatment of *Staphylococcus aureus* infections. (i) This K-like phage has a broad host range; all 83 tested clinical isolates of *S.aureus* tested were susceptible to PYO^Sa^. (ii) Because of the mode of action of PYO^Sa^ *S. aureus* is unlikely to generate classical receptor-site mutants resistant to PYO^Sa^; none were observed in the 13 clinical isolates tested. (iii) PYO^Sa^ kills *S. aureus* at high rates. On the downside, the results of our experiments and tests of the joint action of PYO^Sa^ and antibiotics raise issues that must be addressed before PYO^Sa^ is employed clinically. Despite the maintenance of the phage, PYO^Sa^ does not clear the populations of *S. aureus*. Due to the ascent of a phenotypically diverse array of small colony variants following an initial demise, the bacterial populations return to densities similar to that of phage-free controls. Using a combination of mathematical modeling and *in vitro* experiments, we postulate and present evidence for a mechanism to account for the demise–resurrection dynamics of PYO^Sa^ and *S. aureus*. Critically for phage therapy, our experimental results suggest that treatment with PYO^Sa^ followed by bactericidal antibiotics can clear populations of *S. aureus* more effectively than the antibiotics alone.

**Significance Statement:** The increasing frequency of antibiotic-resistant pathogens has fostered a quest for alternative means to treat bacterial infections. Prominent in this quest is a therapy that predates antibiotics: bacteriophage. This study explores the potential of a phage, PYO^Sa^, for treating *Staphylococcus aureus* infections in combination with antibiotics. On first consideration, this phage, isolated from a commercial therapeutic cocktail, seems ideal for this purpose. The results of this population dynamic and genomic analysis study identify a potential liability of using PYO^Sa^ for therapy. Due to the production of potentially pathogenic atypical small colony variants, PYO^Sa^ alone cannot eliminate *S. aureus* populations. However, we demonstrate that by following the administration of PYO^Sa^ with bactericidal antibiotics, this limitation and potential liability can be addressed.

## Introduction

Driven by well-warranted concerns about the increasing frequencies of infections with antibiotic-resistant pathogens, there has been a resurrection of interest in, research on, and clinical trials with a therapy that predates antibiotics by more than 15 years: bacteriophage (1–4). One direction phage therapy research has taken is to engineer lytic (virulent) phages with properties that are anticipated to maximize their efficacy for treating bacterial infections in mammals (5–7). Primary among these properties are (i) a broad host range for the target bacterial species, (ii) mechanisms that prevent the generation of envelope or other kinds of high-fitness resistance in the target bacteria (8), (iii) the capacity to thwart the innate and adaptive immune systems of bacteria, respectively restriction-modification and CRISPR-Cas (7, 9, 10), (iv) the ability to survive, kill, and replicate on pathogenic bacteria colonizing or infecting mammalian hosts (11, 12) and (v) little or no negative effects on the treated host (8).

To these five desired properties for therapeutic bacteriophage, we add a sixth: synergy with antibiotics. Phage-only treatment may be reasonable for compassionate therapy where the bacteria responsible for the infection are resistant to all available antibiotics (13–15). But from a practical perspective, for phage to become a widely employed mechanism for treating bacterial infections, they would have to be effective in combination with antibiotics. It would be unethical and unacceptable to clinicians and regulatory agencies to use phage independently for infections that can be effectively treated with existing antibiotics.

Although not specifically engineered for these properties, there is a *Staphylococcal* phage isolated from a therapeutic phage collection from the Eliava Institute in Tbilisi; we call PYO^Sa^ that on first consideration appears to have all six of the properties required to be an effective agent for therapy. (i) PYO^Sa^ is likely to have a broad host range for *S. aureus*. The receptor of this K-like Myoviridae is N-acetylglucosamine in the wall-teichoic acid backbone of *Staphylococcus aureus* and is shared among most (16), if not all *S. aureus*; thereby suggesting PYO^Sa^ should be able to adsorb to and potentially replicate on and kill a vast number of clinical isolates of *S. aureus*. (ii) *S. aureus* does not generate classical, surface modification mutants resistant to PYO^Sa^. Since the structure of the receptor of PYO^Sa^ is critical to the viability, replication, and virulence of these bacteria, the modifications in this receptor (17) may not be consistent with the viability or pathogenicity of *S. aureus* (18). (iii) The replication of PYO^Sa^ is unlikely to be prevented by restriction-modification (RM) or CRISPR-Cas. Despite a genome size of 127 KB, the PYO^Sa^ phage has no GATC restriction sites for the *S. aureus* restriction enzyme Sau3A1 and only one restriction site, GGNCC, for the Sau961 restriction endonuclease (19, 20). There is no evidence for a functional CRISPR-Cas system in *S. aureus* or, to our knowledge, other mechanisms by which *S. aureus* may prevent the replication of this phage (21). (iv) There is evidence that PYO^Sa^–like phages can replicate in mammals. Early treatment with a phage with a different name but the same properties as PYO^Sa^, Statu^v^, prevented mortality in otherwise lethal peritoneal infections of *S. aureus* in mice (22). A PYO^Sa^-like phage has also been successfully used therapeutically in humans (23). (v) No deleterious effects of a PYO^Sa^-like phage were observed in recent placebo-controlled trials with volunteers asymptotically colonized by *S. aureus* (19). (vi) Finally, there is evidence to suggest synergy with antibiotics. *In vitro*, PYO^Sa^ increased the efficacy of low concentrations of antibiotics for the treatment of biofilm populations of *S. aureus* (24).

With *in vitro* parameter estimation, population and evolutionary dynamic studies, and experiments with PYO^Sa^ and *S. aureus* Newman in combination with three different bacteriostatic and six different bactericidal antibiotics, we explore just how well PYO^Sa^ fits the above criteria for combination antibiotic and phage therapy. Our results suggest that PYO^Sa^ scores well on most of these tests but does not get an “A”. As a consequence of selection for potentially pathogenic small colony variants, by itself PYO^Sa^ does not clear *S. aureus* infections; after an initial demise when confronted with this phage, although the phage continue to be present, the densities of the bacteria return to levels similar to those observed in the absence of this virus. By employing antibiotics and phage against *S. aureus* we could demonstrate synergy in clearing the bacterial population. However, there were significant differences in effectiveness depending on whether the antibiotics and phage were used simultaneously or in succession, and on whether the antibiotics used were bacteriostatic or bacteriocidal. Our most important result from a therapeutic perspective is that treatment of *S. aureus* cultures with PYO^Sa^, followed by the administration of bactericidal antibiotics, is more effective at clearing the bacterial population than treatment with these antibiotics alone.

## Results

### (1) Bacteriophage PYO^Sa^ has a broad host range for *S. aureus*

We use two assays to determine the host range of PYO^Sa^: (i) the production of zones of inhibition in soft agar lawns and (ii) changes in the optical density of exponentially growing liquid cultures of *S. aureus* mixed with this phage. By both criteria, *S. aureus* Newman and 12 clinical isolates of methicillin-sensitive *S. aureus* from the Network on Antimicrobial Resistance (NARSA) collection (25) were all sensitive to PYO^Sa^ and appeared to be unable to generate classically resistant mutants. Additional evidence for a broad host range of PYO^Sa^ comes from a survey of 71 clinical isolates of *S. aureus*, including 54 MRSA strains and phylogenetically similar species performed by LabCorp™ (*SI Appendix*, Table S1).

### (2) *S. aureus* appears to be unable to generate classical, surface modification, mutants resistant to PYO^Sa^

Evidence for this comes from experiments with *S. aureus* Newman and 12 clinical isolates from the NARSA collection (25). Single colonies of each strain were grown in the presence of ~10^6^ phage particles per ml, and after 24 hours of exposure to the phage in liquid culture, the optical densities of exponentially growing *S. aureus* were no greater than that of the media without the bacteria, and by plating, we were unable to detect colonies from these cultures.

### (3) Bacterial population heterogeneity and the maintenance of phage

With the bacterial growth and phage infection parameters estimated for *S. aureus* Newman and PYO^Sa^, a complete clearance of the bacteria in these experiments after less than 24 hours of exposure to PYO^Sa^ is anticipated from a simple mass action model of the population dynamics of bacteria and phage (26). To determine whether this is the case empirically, we divided a culture with ~ 10^4^ *S. aureus* Newman into 28 tubes. We let these cultures replicate for 2 hours, for an average density of 3 × 10^7^ and then added ~6 × 10^8^ PYO^Sa^. At 24 hours, all 28 cultures were clear, and no bacterial colonies were found on the LB plate samplings of these cultures. However, by day seven, 26 of these 28 independent cultures were turbid or somewhat turbid. These turbid cultures all had phage at densities in excess of 5×10^6^. Upon plating these 7-day cultures, colonies were detected with two distinct phenotypes: colonies similar in size to the ancestral wild-type *S. aureus* Newman and much smaller colonies, small colony variants (Fig. 1*A*).

**Fig. 1.**
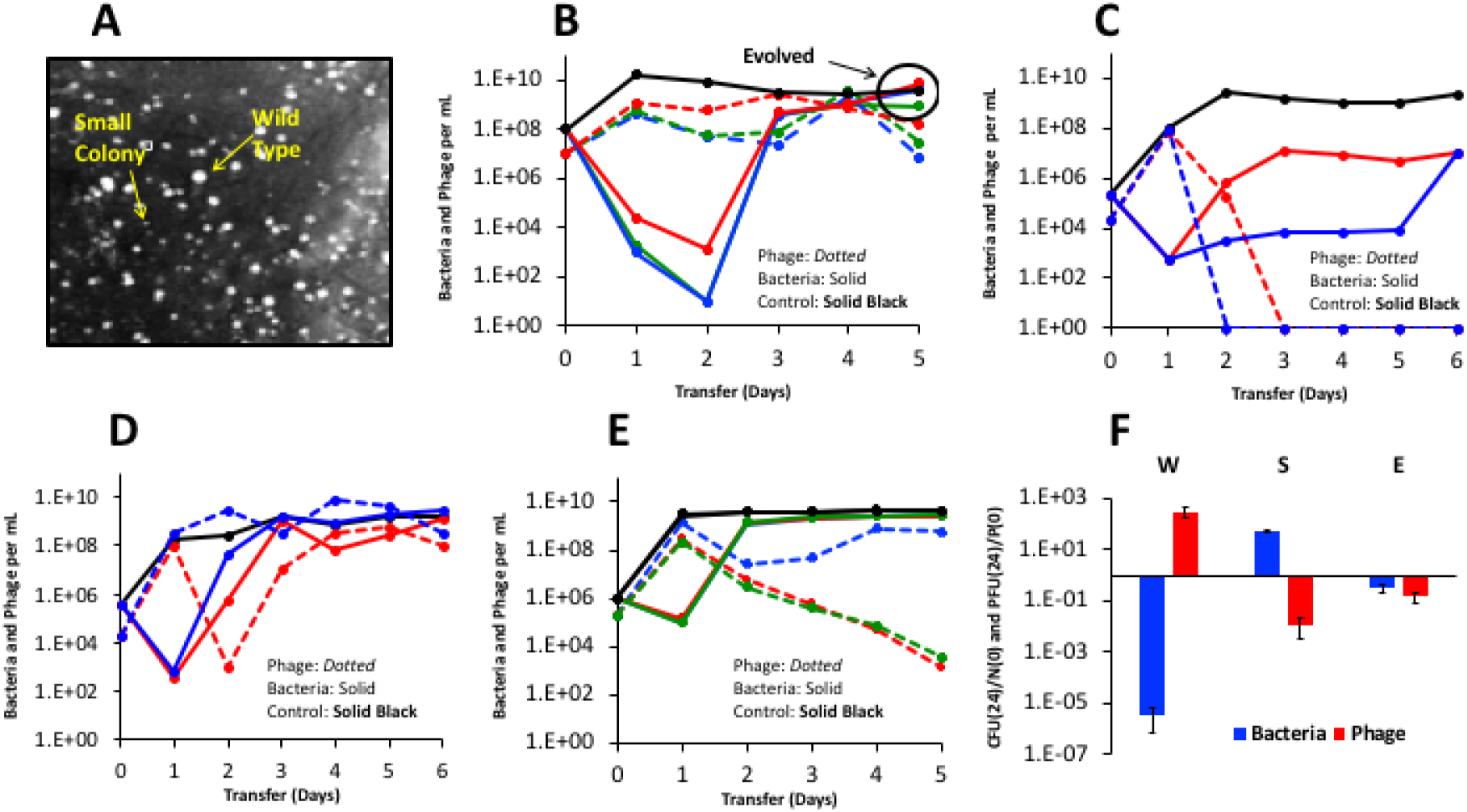
Small colony variants and the population dynamics of PYO^Sa^ and *S. aureus* Newman in serial transfer culture. **A.** Wild type *S. aureus* and small colony variants visualized. **B.** Changes in densities of *S. aureus* Newman and PYO^Sa^ in three independent (red, green, blue) serial transfer experiments diluted 1/100 in fresh media daily. **C.** Changes in densities of *S. aureus* Newman and PYO^Sa^ in serial transfer cultures initiated two small colony variants isolated from the 5^th^ transfer of the cultures with PYO^Sa^ and *S. aureus* Newman in panel B. **D.** Changes in densities of *S. aureus* Newman and PYO^Sa^ in serial transfer cultures initiated with PYO^Sa^ and equal densities of cultures derived from small colonies and the ancestral *S. aureus* Newman. **E.** Changes in densities of *S. aureus* and PYO^Sa^ in serial transfer cultures initiated with PYO^Sa^ and single colonies of evolved (wild-type colony morphology) bacteria isolated from the 5^th^ transfer of the cultures in panel B. **F.** Changes in the ratio of bacteria and phage at time 0 and 24 hours. Wild type *S. aureus* Newman (W), small colony variants (S), and evolved bacteria (E). Three independent replicas.

To elucidate why PYO^Sa^ does not kill all of the *S. aureus* Newman in these liquid cultures, we prepared three independent 10 ml cultures with ~ 10^8^ *S. aureus* Newman and ~10^7^ PYO^Sa^ and serially passaged these cultures for five days, transferring 1/100 of these cultures to fresh media each day. The results of these serial transfer experiments are presented in Fig. 1*B*. As anticipated from the experiment with the 28 tube cultures, at 24 hours, all the flasks were clear. The densities of bacteria remaining in these cultures estimated by plating, ~10^3^ per ml, is probably an underestimate of the real density due to killing by phage on the plates. Additional support for the hypothesis that the densities of *S. aureus* in these liquid cultures is substantially greater than the CFU estimate is that the density of the PYO^Sa^ phage did not decline at each transfer, and therefore, the phage must have been replicating. This hypothesis was evaluated with a simple model (26) using the phage infection parameters estimated for PYO^Sa^ and *S. aureus* Newman. To maintain a population in a culture diluted by 100-fold the phage density must increase by at least 100-fold at each transfer and for this to occur, the product of the adsorption rated constant, δ, the burst size β, and the density of sensitive bacteria, N, has to exceed 100. With the estimated values, δ=4.8×10^−7^ and β=80, the density of bacteria in these cultures would have to be at least 2.6×10^6^ for the phage not to be diluted out.

Most intriguingly, in all three independent serial transfer populations with the PYO^Sa^ phage present, the CFU estimates of the density of *S. aureus* started to increase after the 2^nd^ transfer. By the 4^th^ and 5^th^ transfer, the density of cells in these cultures was similar to that of the phage-free controls, and the phage continued to be maintained, presumably because of the continued presence of phage-susceptible populations. To test this we spread bacterial aliquots onto agar and observed heterogeneity in colony growth with both fast-growing and small colony variants. Using a spot test we showed that cultures grown from fast-growing colonies, henceforth called the evolved bacteria, were susceptible to the original PYO^Sa^ as well as to the co-existing PYO^Sa^. In contrast, the spot test showed that cultures grown from small colony variants isolated from these late transfer cultures were resistant to lysis by PYO^Sa^.

To further evaluate the different bacterial phenotypes present in the late transfer cultures we serially passaged two small colony cultures in the presence of PYO^Sa^ (Fig. 1*C*). We observed that cultures generated from small colony variants were unable to support the phage. These data, and the spot test data suggest an explanation for the maintenance of bacteria and phage throughout the original transfer experiment (Fig. 1*B*). Our hypothesis is that the bacterial population is polymorphic and includes sensitive cells capable of supporting phage growth as well as a non-sensitive population. We tested this hypothesis by mixing wild-type *S. aureus* Newman in equal frequency with small colony variants, and observed, consistent with the hypothesis, that the phage were maintained throughout the serial transfers and that the total bacterial population density remained similar to that in phage-free controls (Fig. 1*D*). As a separate test of the polymorphic population hypothesis, we made serial transfer experiments initiated with mixtures of PYO^Sa^ phage and evolved bacteria isolated from the 5^th^ transfer cultures (Fig. 1*E*). The results of these experiments are consistent with this reduced susceptibility hypothesis. Moreover, within a single transfer, the bacterial density returned to that of the phage-free controls and the phage declined, albeit at a rate less than that anticipated if they were not replicating at all, in two of the three cultures. In the two cultures where the phage declined significantly, the dominant bacterial population were small colony variants; while, the replicate with the co-existing phage was dominanted by bacteria that generated wild-type colony morphologies. See *SI Appendix*, Section II for a consideration of the between experiment variation in the population dynamics observed in Fig. 1*B* and 1*E*.

Based on the preceding results and interpretations, we postulate that the bacteria are of three states, the ancestral wild-type, small colony variants, and what we call the evolved state, which have wild-type or near wild-type colony growth rate and rise to high densities after serial passaging in the presence of phage. Both the wild-type and the evolved are postulated to be capable of supporting phage growth whereas the small colonies type do not support phage growth.

### (4) The genetic basis of bacterial colony growth rate variation

To determine the genetic basis of the differences between wild-type *S. aureus* Newman, the small colony variants, and evolved states, colonies with a range of sizes were isolated from early and late in serial passaging experiments similar to those shown in Fig. 1*B*. Whole genome sequencing of several clones revealed a clear sequence of events during the passaging. The first event was the selection of mutations in *femA* which encodes a protein responsible for assembling the pentaglycine interpeptide bridges in the *S. aureus* cell wall (*SI Appendix*, Table S2). These mutants all have a small colony variant phenotype. The range of observed colony sizes is representative of the diversity of mutations identified in *femA*, and presumably indicates a direct correlation between the growth rate and the severity of the defect in FemA function. We observed in later transfers larger colonies that carried a *femA* mutation and an additional mutation (see below) suggesting that this was a compensatory mutation restoring growth rate and creating the evolved state. The small colony variant phenotype of *S. aureus* is known to be very unstable and subject to rapid suppression by a wide variety of compensatory mutations (27). To investigate the nature of evolved state, we chose several different *femA* mutants and used these to select faster-growing colonies. Fast-growers were easily selected and were observed on agar as larger colonies growing above the slower-growing parental small colony strains. Whole genome sequencing revealed an array of suppressing mutations were selected, including an internal suppressor in *femA*, mutations in a variety of genes affecting secondary messenger metabolism, and frequent mutations in the transcriptional regulator *sarA* (*SI Appendix*, Table S3). We concluded that the initial event upon exposure of wild-type *S. aureus* Newman to PYO^Sa^ is the selection of small colony variants associated with mutations in *femA*. These slow-growing mutants then evolve to the faster-growing evolved state by the acquisition of suppressor mutations that frequently affect global transcriptional regulators.

### (5) Phenotypic differences between wild-type, small colony, and evolved clones

The hypothesis above predicts that cultures derived from bacterial colonies of each state should respond differently when exposed to PYO^Sa^. To test this, we co-cultured high densities of each bacterial state (W, S, and E) with PYO^Sa^ and determined the relative change in densities of both bacteria and phage, respectively 10^8^ and 10^7^ cells and particles per ml, after 24 hours exposure (Fig. 1*F*). Each of the three states exhibited a unique dynamic. The density of the wild-type and evolved bacteria declined, with the wild-type declining to a much greater extent than the evolved state. In contrast, the density of small colony variant bacteria increased. PYO^Sa^ phage increased on the wild-type bacteria but declined on both small colony variant and evolved state bacteria. The decline in phage was greater on small colony variants than on the evolved state.

Measurement of the growth rates in liquid culture of the three bacterial states confirmed the growth parameters previously inferred from colony sizes (*SI Appendix*, Table S4). Mutations causing an small colony phenotype on agar also caused severe growth defects in liquid culture, while the evolved clones with secondary suppressing mutations showed partial restoration of growth rate toward wild-type levels.

The pentapeptide crosslinks that FemA synthesizes are notably the target of lysostaphin. FemA mutants resistant to lysostaphin have previously been observed to become hypersensitive to penicillin antibiotics (28, 29). To test for evidence of these phenotypes the MIC of the three states were measured against lysostaphin and oxacillin (*SI Appendix*, Table S4). We observed that all of the strains carrying *femA* mutations (both small colony and evolved state) became resistant to lysostaphin and hypersensitive to oxacillin. Accordingly, the compensatory evolution from small colony variant to evolved state does not phenotypically recreate a wild-type phenotype: wild-type and evolved are genetically and phenotypically distinct states. Since FemA is responsible for adding the 2^nd^ and 3^rd^ glycines to the interpeptide bridge, while FemB is responsible for adding the 4^th^ and 5^th^ residues, mutations in *femA* are expected to result in single glycine bridges between the cell wall peptides. To the degree that the *femA* mutants identified have different growth and susceptibility phenotypes, the cell wall defect in strains carrying these mutations is likely not complete, but rather represents a balance between full length pentapeptide crosslinks and single glycine crosslinks. FemA was recently reported to be essential (30) and it is notable that in our experiments no unequivocally *femA* null mutations were identified. Based on the number of mutants we have screened, this indicates that *femA* null mutants are either non-viable or are counterselected in these experiments.

### (6) PYO^Sa^ does not replicate on stationary phase *S. aureus*

In infected hosts, many of the bacteria will not be replicating, and thereby, phage would be particularly effective for treatment if they, like some antibiotics (31), could kill non-replicating *S. aureus*. To determine whether PYO^Sa^ can kill and replicate on non-growing bacteria, three independent 48-hour stationary phase cultures were mixed with PYO^Sa^ for average initial densities of ~4×10^9^ *S. aureus* and 10^6^ PYO^Sa^. The bacteria and phage were incubated with shaking for 24 hours, and the viable cell and phage densities were estimated and compared to the initial densities. There was no evidence for the stationary phase bacteria being killed, the mean and standard error of the N(24)/N(0) ratio was 0.96 ± 0.03. Moreover and critically, there was a significant decline in the density of phage, N(24)/N(0) = 0.043 ± 0.006. Thus, not only does PYO^Sa^ not kill stationary phase *S. aureus*, these non-replicating bacteria act as a sink and could reduce the density of PYO^Sa^ from treated hosts.

The results of experiments to determine whether the decline in the density of phage can be attributed to changes in the medium, such as a high pH, were negative. The density of PYO^Sa^ did not decline in sterile filtrates of 48-hour stationary phase cultures. There is, however, the suggestion that the bacteria must be viable to lead to the reduction in the viable density. When the *S. aureus* in the 48-hour stationary phase culture are killed with chloroform, the density of PYO^Sa^ does not decline

### (7) Accounting for the population dynamics presented in Fig. 1*B*

In the *SI Appendix*, Section IV, Fig. S2, using a mathematical model and numerical solutions, we present a hypothesis for the kill-recovery dynamics observed for the bacteria, and the maintenance of the phage in the serial transfer cultures depicted in Fig. 1*B*. In accord with this hypothesis, (i) the bacterial population recovers from its initial demise due to predation by PYO^Sa^ phage, by the generation and ascent of small colony variants which are immune to, and selected by, the phage, and (ii) the phage are maintained because of the instability of the small colony variants which continously generate the evolved state bacteria upon which the phage can replicate.

### (8) The joint action of PYO^Sa^ and antibiotics

To determine whether the action of antibiotics and PYO^Sa^ would be synergistic or antagonistic in killing *S. aureus* Newman and the effect of these drugs on replication of PYO^Sa^, we followed the change in the densities of bacteria and phage over 24 hours of exposure to antibiotics and phage in combination, and to phage alone. For these experiments, we mixed growing cultures of *S. aureus* Newman and PYO^Sa^ in MHII at a density of 4×10^6^ with 4×10^5^, respectively and super MIC concentrations of the antibiotics. The densities of bacteria and phage were estimated just before the antibiotics were added and 24 hours later, respectively N(0) and N(24), where N is either CFU or PFU. In these experiments, the antibiotics and phage were introduced into the growing cultures of *S. aureus* Newman in three different ways: (i) simultaneously, AB+PYO^Sa^, (ii) antibiotics first and phage 30 minutes later, AB→PYO^Sa^, and (iii) phage first and antibiotics 30 minutes later, PYO^Sa^→AB. As controls, we also treated parallel cultures of *S. aureus* Newman with antibiotics only, AB, and with phage alone. In Fig. 2 (bacteriostatic antibiotics) and Fig. 3 (bacteriocidal antibiotics) we present N(24)/N(0) ratios for the bacteria and phage for three independent experiments with each antibiotic and phage combination.

**Fig. 2.**
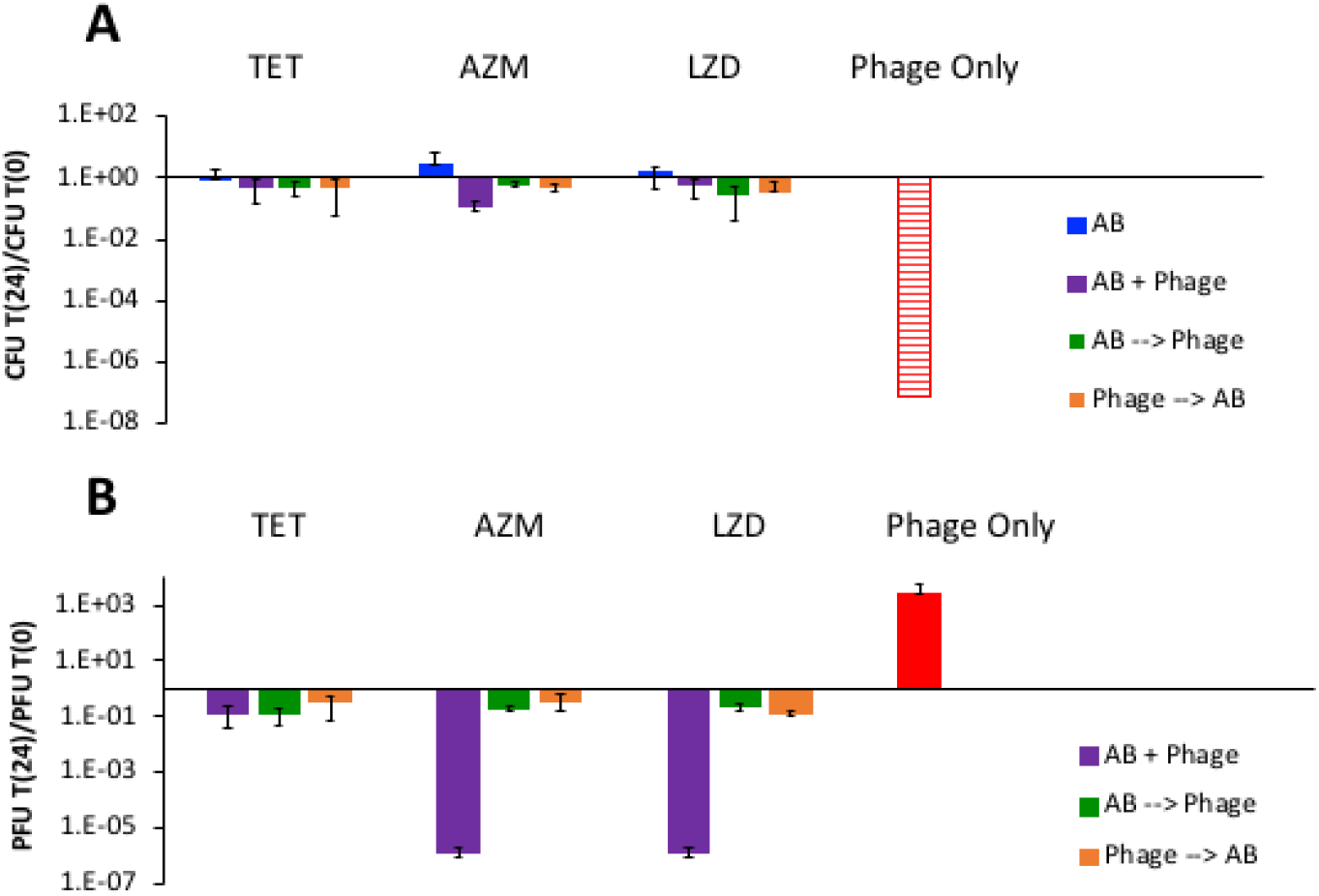
Joint action of bacteriostatic antibiotics and PYO^Sa^. The concentrations of these antibiotics are 10μg/ml. **A.** The ratio of the change in density of *S. aureus* after 24 hours of exposure to antibiotics (blue), antibiotics plus phage (purple, green, orange, refers to order of addition as explained in the text) or phage alone (red). Hash red, the density of *S. aureus* recovered was below the detection limit, ~10^2^ cells per ml. **B.** The ratio of the change in the density of PYO^Sa^ after 24 hours of confronting wild-type *S. aureus* in combination with antibiotics or alone.

**Fig. 3.**
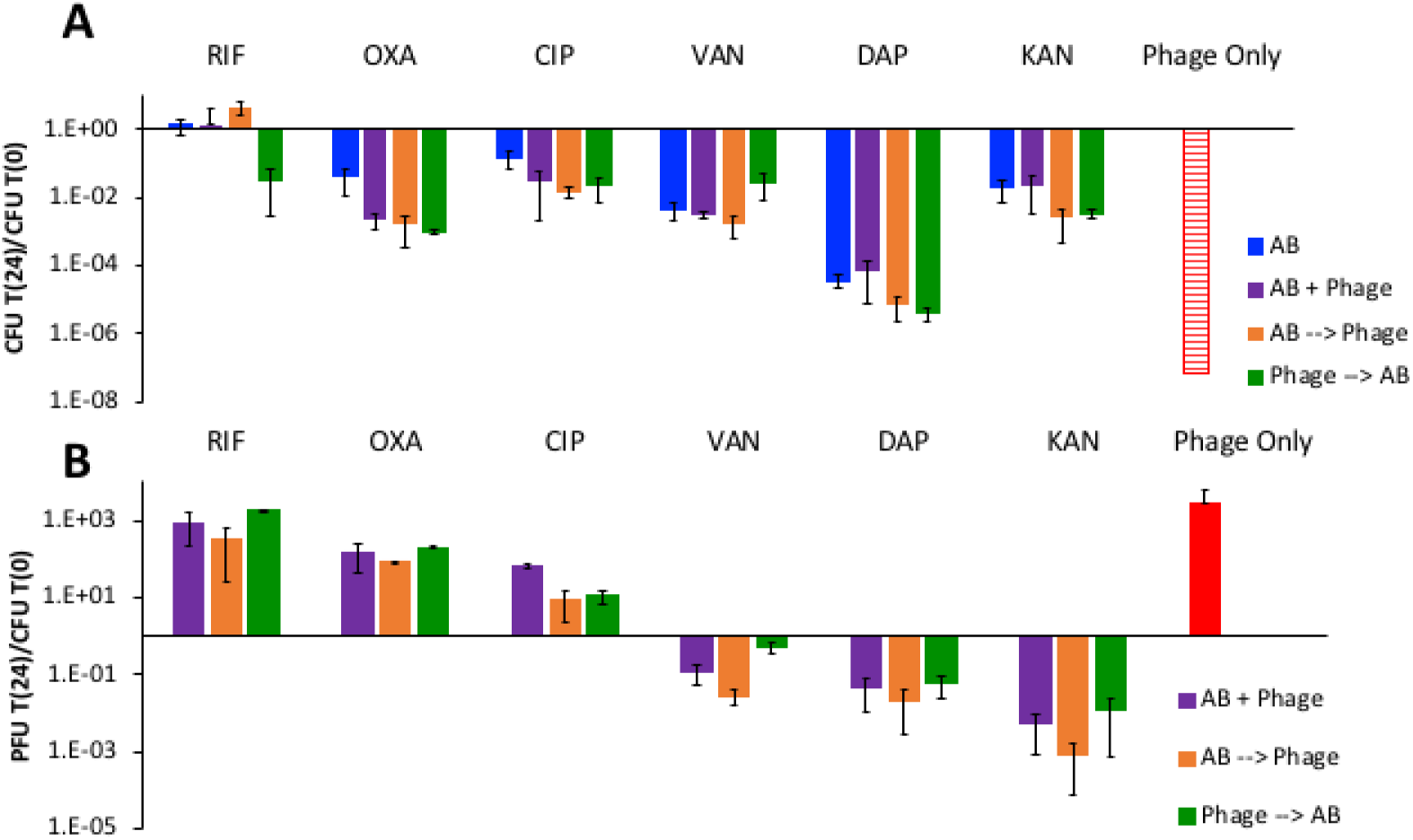
The joint action of bactericidal antibiotics and PYO^Sa^. Concentrations of the different antibiotics in μg/ml (RIF 0.02, OXA 3, CIP 0.5, VAN 8, DAP 64, KAN 46), corresponding to minimum bacteriocidal concentrations. **A.** The ratio of the change in density of *S. aureus* after 24 hours of exposure to antibiotics, antibiotics and phage, or phage alone. Hash red, the density of *S. aureus* recovered was below the detection limit, ~10^2^ per ml. **B.** The ratio of the change in the density of PYO^Sa^ after 24 hours of confronting *S. aureus* Newman in combination with antibiotics or alone.

With the bacteriostatic antibiotics, tetracycline (TET), azithromycin (AZM), and linezolid (LZD), the greatest decline in the density of bacteria and increase in the density of phage obtained in the experiment where phage are used alone. There is clearly a negative synergy between these bacteriostatic antibiotics and the phage. Whether administered simultaneously or sequentially, at the concentrations used, these antibiotics prevent PYO^Sa^ from killing *S. aureus* Newman. When the phage are administered simultaneously with azithromycin and linezolid, the phage density declines.

In Fig. 3, we present the results of experiments with bactericidal antibiotics, rifampin (RIF), oxacillin (OXA), ciprofloxacin (CIP), vancomycin (VAN), daptomycin (DAP) and kanamycin (KAN). The data suggest that the simultaneous or sequential administration of PYO^Sa^ may modestly increase the rate at which bacteriocidal antibiotics kill *S. aureus* at the concentrations employed. However, as was observed for the parallel experiment with the bacteriostatic drugs (Fig. 2), the phage kill more *S. aureus* in the absence of antibiotics than with these drugs, an antagonistic interaction once again. The failure of rifampin to reduce the viable density of *S. aureus* can be attributed to rifampin resistance emerging. It should be noted however, that in one of the three treatments where PYO^Sa^ was used before adding rifampin, it prevented the ascent of resistance. Most interestingly, while treatment with rifampin, oxacillin, and ciprofloxacin allowed PYO^Sa^ to replicate, this was not the case for vancomycin, daptomycin, and kanamycin, which appeared to suppress the replication of PYO^Sa^.

### (9) Sequential treatment, phage followed by antibiotics

The serial transfer results presented in Fig. 1*B* indicate that as a consequence of the emergence and ascent of small colony variants, PYO^Sa^ by itself will not be able to control an *S. aureus* population for an extended time. Although the phage continue to be present, the density of the bacteria returns to levels similar to that of the phage-free control (Fig. 1*B*). We asked what result would be obtained if antibiotics were administered to these populations? To address this question, we performed the serial transfer experiments with the addition of bactericidal antibiotics following the phage-mediated reduction in the density of wild-type *S. aureus* observed during the first 24 hours. The results of these experiments are presented in Fig. 4.

**Fig. 4.**
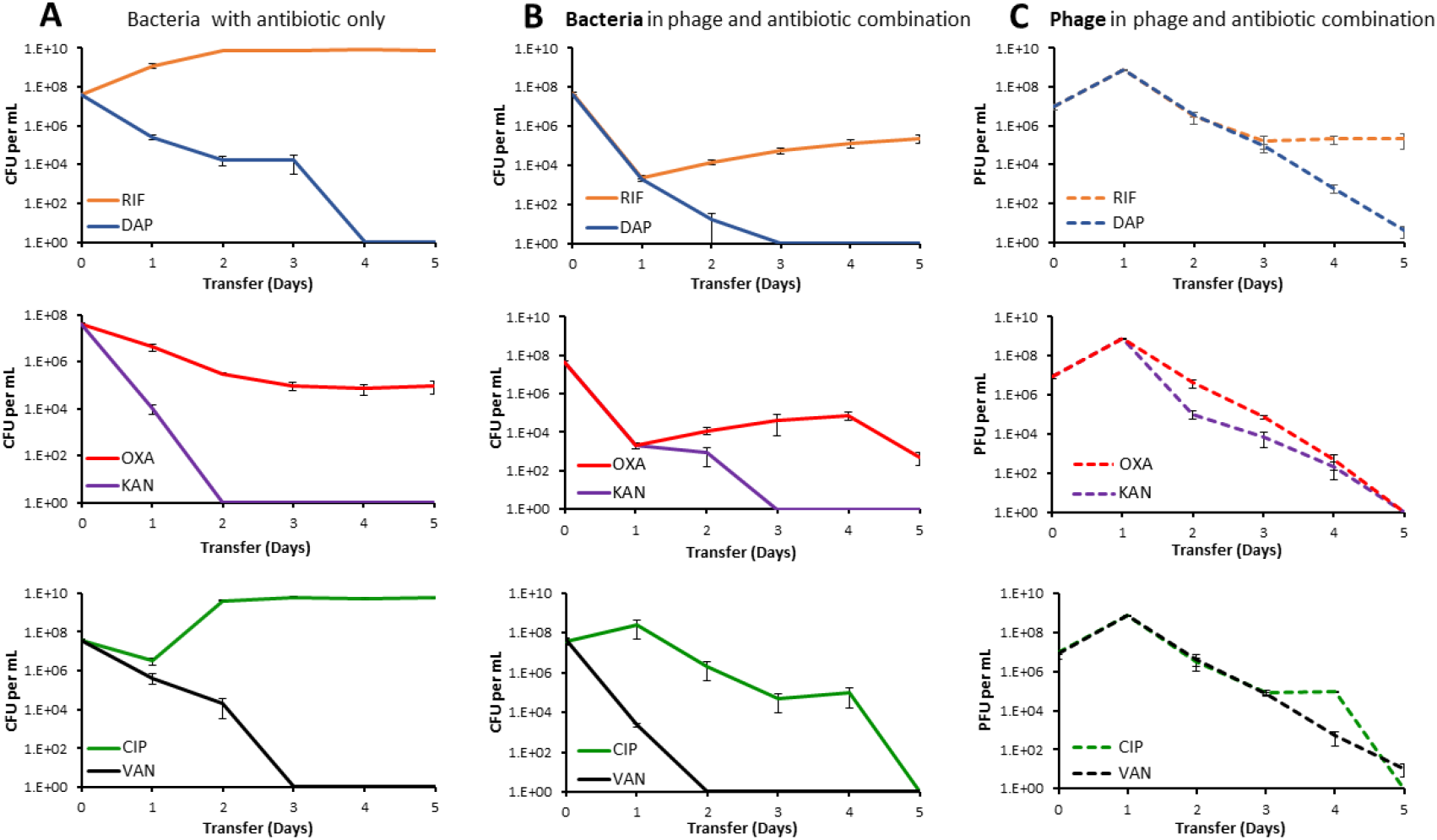
Changes in the densities of bacteria and phage in serial transfer cultures treated with antibiotics alone, or in combination with phage. **Column A.** Bacteria treated with antibiotics alone. **Column B.** Bacterial densities in cultures containing both the antibiotic and the phage. **Column C.** Phage densities in cultures containing both the antibiotic and the phage. Mean and standard errors of 3 independent experiments. **Row 1.** Treatment with RIF and DAP, 0.02 and 64 μg/ml, respectively. **Row 2.** Treatment with KAN and OXA, 46 and 3 μg/ml respectively, **Row 3.** Treatment with CIP and VAN, 0.5 and 8 μg/ml respectively.

Three of the antibiotics, DAP, KAN and VAN, were alone sufficient to eliminate the *S. aureus* population (Fig. 4*A*). In contrast, with RIF by the second transfer the density of *S. aureus* in treated cultures reached a density observed for antibiotic and phage free cultures (~2×10^9^ bacterial per ml). The reason for the failure of RIF to clear the culture was the emergence of mutants resistant to this drug. All six colonies tested from the 5^th^ transfer were resistant to RIF. However, the viable cells recovered from the phage and RIF combination cultures were as sensitive to RIF as the antibiotic-free controls. Neither OXA nor CIP alone cleared the cultures. In the case of OXA, the viable cell density declined but continued to persist at a density of approximately 10^5^ cells per ml. The bacteria recovered from these cultures were sensitive to OXA. We postulate that this leveling off can be attributed to persistence (32); see *SI Appendix*, Section VI, Fig. S5. In the case of CIP, the initial exposure led to a substantial decline in the viable cell density of *S. aureus*, however, following the second transfer the bacterial population recovered and was sustained at densities similar to that in the antibiotic-free controls. The colonies of *S. aureus* recovered at the end of this experiment were susceptible to CIP. We postulate that these resurrection dynamics could be attributed to heteroresistance (33), see *SI Appendix*, Section VI, Fig S5.

## Discussion and conclusions

On first consideration, PYO^Sa^ seems to be an ideal phage for treating *S. aureus* infections. The results of this study provide evidence in support of three virtues of PYO^Sa^ as a therapeutic phage. 1) PYO^Sa^ is likely to kill virtually all methicillin-resistant as well as methicillin-sensitive *S. aureus*. 2) *S. aureus* are unable to generate classical surface-resistant mutants to PYO^Sa^,^,^ thus, cocktails of multiple phages would not be needed to ensure coverage or prevent resistance. 3) *S. aureus* Newman has a high adsorption rate and burst size with PYO^Sa^ and, when first confronting growing populations of *S. aureus*, the bacteria are killed, and the phage replicate at a high rate.

On the downside, our experiments raise caveats about the use of PYO^Sa^ for treating *S. aureus* infections alone and suggest a possible liability as well. Not only is PYO^Sa^ unable to clear cultures of *S. aureus* Newman, it selects for potentially pathogenic small colony variants, (34–37) that are refractory to this phage. Through a “leaky resistance” mechanism (38), the phage continue to be maintained, and the bacterial population continues to persist at densities not much less than they do in the absence of PYO^Sa^. Although not observed for the small colony variants tested here, at least some small colonies are more resistant to antibiotics than the bacteria from which they are derived (27, 39, 40).

For phage therapy to be a practical and acceptable enterprise, these viruses would have to be used in combination with antibiotics. Our results indicate that when administered simultaneously, or nearly simultaneously with antibiotics, PYO^Sa^ does worse in killing *S. aureus* than it does alone PYO^Sa^. The ribosome-targeting bacteriostatic antibiotics, tetracycline, azithromycin, and linezolid, suppress the ability of PYO^Sa^ to kill *S. aureus* Newman. This observation is consistent with the failure of PYO^Sa^ to replicate on stationary phase populations of *S. aureus*. It is also consistent with the Numbers Game hypothesis for the action of bacteriostatic antibiotics (41) according to which, the number of free ribosomes is too low to support the protein synthesis needed for replication of the phage.

When administered simultaneously with bactericidal antibiotics, PYO^Sa^ is also less effective in killing *S. aureus* than it is in the absence of these drugs. We postulate that this can be attributed to pharmaco and population dynamics of the joint action of antibiotics and phage; the antibiotics reduce the densities of the bacteria substantially and thereby lower the capacity of the phage to replicate (*SI Appendix*, Section IV).

### Sequential treatment with PYO^Sa^ and antibiotics

Our experiments suggest a way to deal with the major caveat and potential liability of treatment with PYO^Sa^ alone, the recovery of the bacterial population due to the ascent of small colony variants. The administration of bactericidal antibiotics following the initial decline in the density of bacteria due to PYO^Sa^ prevents bacterial population recovery and eliminates or prevents the selection of small colony variants. The latter is not the case when the antibiotics are used alone. One interpretation of this is that sequential treatment, initially with phage, then with a bactericidal antibiotic may be more effective than treatment with antibiotics alone or phage alone.

### Conclusion and Recommendation

We interpret the results of this in silico and *in vitro* study to suggest that PYO^Sa^ will be effective for treatment of *S. aureus* infections, but only if the administration of bactericidal antibiotics follows that of phage. The next step will, of course, be to test this sequential phage and antibiotic treatment hypothesis with *S. aureus* infections in experimental animals.

We suggest that the *in vitro* methods used to explore the potential efficacy of PYO^Sa^ would also be useful for evaluating other phages being developed for treating bacterial infections. Existing data suggested that PYO^Sa^ met all of the criteria desired for a phage to be effective for therapy; yet, the *in vitro* experiments performed here uncovered a limitation and potential liability of using this phage for therapy that would not have anticipated and they also subsequently revealed a way to deal with said limitation and liability.

## Materials and Methods

### Strains and growth media

Unless otherwise noted, all experiments were performed from derivatives of the parent strain *S. aureus* Newman (ATCC 25904). The parent *S. aureus* Newman was obtained from Bill Schafer of Emory University. The small colony variant and evolved strains were obtained from PYO^Sa^ challenged *S. aureus* Newman by experiments performed in our lab. The investigation for classical resistance was performed in the following MSSA strains obtained from Abraham Moller in the Reid Lab at Emory University: NRS52, NRS102, NRS180, NRS110, NRS252, NRS253, NRS266, NRS109, NRS148, and NRS205.

Bacterial cultures were grown at 37°C in Mueller-Hinter II (MHII) Broth [275710, BD™] and on Luria-Bertani Agar (LB) Plates [244510, BD™]. PYO^Sa^ lysates were prepared from single plaques at 37°C in MHII broth alongside wild-type *S. aureus* Newman by plate lysis. Specifically, individual phage plaques were picked with a sterile stick, resuspended in 4ml of soft agar with 0.1ml of overnight bacterial culture and plated on top of phage plates. The plates were then incubated at 37°C overnight. The soft agar was scraped with a sterile iron scoop, resuspended in 10ml MHII with ~0.5ml of chloroform to kill the surviving bacteria. The lysates were then centrifuged to remove the agar, sterilized by filtration (0.2 μm) and stored at 4°C.

### Sampling bacterial and phage densities

Bacteria and phage densities were estimated by serial dilutions in 0.85% NaCl solution followed by plating. The total density of bacteria was estimated on LB (1.6%) agar plates. To estimate the densities of free phage, chloroform was added to suspensions before serial dilutions. These suspensions were mixed with 0.1mL of overnight MHII grown cultures of wild-type *S. aureus* Newman (about 5×10^8^cells per mL) in 4 mL of LB soft (0.65%) agar and poured onto semi-hard (1%) LB agar plates.

### Parameter estimations

The parameters critical for the interaction of the PYO^Sa^ phage and *S. aureus* Newman used in this study were estimated in independent experiments MHII broth. The maximum growth rate of different clones of *S. aureus* Newman was measured by Bioscreen, as described in (42). Phage burst sizes (β) were estimated with one-step growth experiments similar to (43). Adsorption of PYO^Sa^ to *S. aureus* was estimated as described in (43).

### Serial transfer experiments

All serial transfer experiments were carried out in 10ml MHII cultures grown at 37 ̊C with vigorous shaking. The cultures were initiated by 1:100 dilution from 10-mL overnight cultures grown from single colonies. Phage was added to these cultures to reach the initial density of approximately 10^6^ PFU/mL. At the end of each transfer, 0.1mL of each culture was transferred into flasks with fresh medium (1:100 dilution). Simultaneously, 0.1mL samples were taken for estimating the densities of colony-forming units (CFU) and plaque-forming units (PFU), by serial dilution and plating on solid agar.

### Antibiotics and their Sources

Tetracycline, Oxacillin, Vancomycin, Kanamycin, Streptomycin (all from Sigma Aldrich), Azithromycin (Tocris), Daptomycin (MP Biochemicals), Rifampin (Applichem) and Linezolid (Chem-Impex International).

### Whole-genome sequencing

For sequencing individual clones of *S. aureus*, genomic DNA was prepared using the MasterPure Gram-Positive kit, following the manufacturer’s instructions (Epicentre, Illumina Inc., Madison, Wisconsin). Final DNA was resuspended in EB buffer. Genomic DNA concentrations were measured in a Qubit 2.0 Fluorometer (Invitrogen via ThermoFisher Scientific). DNA was diluted to 0.2 ng/μL in water (Sigma-Aldrich, Sweden), and the samples were prepared for whole genome sequencing according to Nextera® XT DNA Library Preparation Guide (Illumina Inc., Madison, Wisconsin). After the PCR clean up-step, samples were validated for DNA fragment size distribution using the Agilent High Sensitivity D1000 ScreenTape System (Agilent Technologies, Santa Clara, California). Sequencing was performed using a MiSeq™ desktop sequencer, according to the manufacturer’s instructions (Illumina Inc., Madison, Wisconsin). The sequencing data were aligned and analyzed in CLC Genomics Workbench version 11.0 (CLCbio, Qiagen, Denmark).

## Acknowledgements

We thank Melony Ivey and Esther Lee for superb technical help. We are grateful to our Staphyloccus maven, Abraham (Jon) Moller, for providing strains and sage advice. And, to Waqas Chaudhry, Andrew Smith, and the reviewers of an earlier incarnation of this report for helpful comments and suggestions. Funds for this research were provided by grants from the US National Institutes of General Medical Sciences, R01 GM091875 and R35 GM 136407 (Bruce Levin) and Vetenskapsrådet (the Swedish Research Council) (2017-03593) and from the Scandinavian Society for Antimicrobial Chemotherapy (SLS-693211, SLS-876451), Diarmaid Hughes. These funders had no role in study design, data collection and interpretation, or the decision to submit the work for publication.

## SUPPLEMENTARY INFORMATION

### I. Host range

In our report, we summarized the evidence that PYO^Sa^’s host range included all 83 clinical isolates of *Staphylococcus aureus* tested. Much of those data are from LabCorp. The methods used are summarized in the following and the results obtained are summarized in Supplemental Table S1.

#### Methods

The methods LabCorp employed, as detailed below, to test the sensitivity of these strains to PYO^Sa^ were somewhat different than those we employed and presented in our report (See Results Section 1).

i. The *S. aureus* strains tested were streaked for single colonies on TSB agar.
ii. These single colonies were inoculated in 3 ml LB broth and grown overnight with shaking at 37°C.
iii. The overnight cultures were diluted by 1:30 in 3ml LB broth and grown with shaking at 37°C to an OD 600nm of between 0.3-1.0.
iv. 300ul of these cultures were added to 3ml LB soft (0.3% or 0.5%) agar supplemented with 3ul 1M CaCl_2_ and 3ul 1M MgCl_2_ which were spread onto LB agar plates to form a lawn.
v. 5μl of the PYO^Sa^ lysates were spotted onto the lawns and incubated overnight.
vi. Lysates were prepared from original spot plates and then re-spotted on the original spot plate host bacteria following the above conditions.

The Key to scoring the results of this assay follows:

| Key |  |
| --- | --- |
| - | No lysis |
| -/+ | Weak lysis |
| + | Partial lysis (cloudy) |
| ++ | Complete lysis |

**Table S1.** Assay for the susceptibility of *S. aureus* to phage PYO^Sa^. Data provided by LabCorp.

| Species | Strain | Spot | Re-spot |
| --- | --- | --- | --- |
| *Staphylococcus aureus* | 27660 | ++ | ++ |
| *Staphylococcus aureus* | RN6390 | + | -/+ |
| *Staphylococcus aureus* | 502A | ++ | ++ |
| *Staphylococcus aureus* | 6538 | ++ | ++ |
| *Staphylococcus aureus* | 25923 | ++ | ++ |
| *Staphylococcus aureus* subsp. aureus | 12600 | ++ | ++ |
| *Staphylococcus aureus* | RN4220 | ++ | ++ |
| *Staphylococcus aureus* subsp. aureus | BAA-1721 | ++ | ++ |
| *Staphylococcus aureus* subsp. aureus | 14775 | ++ | ++ |
| *Staphylococcus aureus* | AR0461 | ++ | ++ |
| Methicillin-resistant *S. aureus* | 017-243 | ++ | ++ |
| Methicillin-resistant *S. aureus* | 041-332 | + | -/+ |
| Methicillin-resistant *S. aureus* | 045-188 | ++ | ++ |
| Methicillin-resistant *S. aureus* | 038-401 | + | ++ |
| Methicillin-resistant *S. aureus* | 049-841 | ++ | ++ |
| Methicillin-resistant *S. aureus* | USA300 | + | - |
| Methicillin-resistant *S. aureus* | 002-639 | + | + |
| Methicillin-resistant *S. aureus* | 049-60 | ++ | ++ |
| Methicillin-resistant *S. aureus* | 026-485 | ++ | ++ |
| Methicillin-resistant *S. aureus* | 011-719 | ++ | ++ |
| Methicillin-resistant *S. aureus* | 045-696 | ++ | ++ |
| Methicillin-resistant *S. aureus* | BAA-1707 | + | -/+ |
| Methicillin-resistant *S. aureus* | BAA-1717 | + | + |
| Methicillin-resistant *S. aureus* | BAA-1720 | ++ | ++ |
| Methicillin-resistant *S. aureus* | BAA-1747 | + | - |
| Methicillin-resistant *S. aureus* | BAA-1754 | ++ | + |
| Methicillin-resistant *S. aureus* | BAA-1761 | ++ | ++ |
| Methicillin-resistant *S. aureus* | BAA-1763 | ++ | ++ |
| **Species** | **Strain** | **Spot** | **Re-spot** |
| Methicillin-resistant *S. aureus* | BAA-1764 | ++ | -/+ |
| Methicillin-resistant *S. aureus* | BAA-1766 | ++ | + |
| Methicillin-resistant *S. aureus* | BAA-1768 | ++ | ++ |
| Methicillin-resistant *S. aureus* | BAA-41 | ++ | ++ |
| Methicillin-resistant *S. aureus* | BAA-42 | ++ | -/+ |
| Multidrug-resistant resistant *S. aureus* | BAA-44 | ++ | + |
| Methicillin-resistant *S. aureus* | BAA-1683 | ++ | + |
| Methicillin-resistant *S. aureus* | BAA-2094 | ++ | ++ |
| Methicillin-resistant *S. aureus* | BAA-2313 | ++ | ++ |
| Methicillin-resistant *S. aureus* | 33592 | ++ | + |
| Methicillin-resistant *S. epidermis* | 700583 | -/+ | - |
| Methicillin-resistant *S. haemolyticus* | 700564 | -/+ | - |
| *Staphylococcus aureus* subsp. aureus | BAA-1718 | ++ | ++ |
| Methicillin-resistant *S. aureus* | AR0462 | ++ | - |
| Methicillin-resistant *S. aureus* | AR0463 | ++ | ++ |
| Methicillin-resistant *S. aureus* | AR0464 | -/+ | -/+ |
| Methicillin-resistant *S. aureus* | AR0465 | ++ | + |
| Methicillin-resistant *S. aureus* | AR0466 | + | - |
| Methicillin-resistant *S. aureus* | AR0467 | ++ | + |
| Methicillin-resistant *S. aureus* | AR0468 | ++ | + |
| Methicillin-resistant *S. aureus* | AR0469 | ++ | + |
| Methicillin-resistant *S. aureus* | AR0470 | ++ | + |
| Methicillin-resistant *S. aureus* | AR0471 | ++ | - |
| Methicillin-resistant *S. aureus* | AR0472 | ++ | ++ |
| Methicillin-resistant *S. aureus* | AR0473 | ++ | ++ |
| Methicillin-resistant *S. aureus* | AR0474 | ++ | + |
| Methicillin-resistant *S. aureus* | AR0475 | + | - |
| Methicillin-resistant *S. aureus* | AR0476 | ++ | ++ |
| Methicillin-resistant *S. aureus* | AR0477 | ++ | ++ |
| Methicillin-resistant *S. aureus* | AR0478 | ++ | ++ |
| Methicillin-resistant *S. aureus* | AR0479 | ++ | ++ |
| **Species** | **Strain** | **Spot** | **Re-spot** |
| Methicillin-resistant *S. aureus* | AR0480 | + | + |
| Methicillin-resistant *S. aureus* | AR0481 | ++ | ++ |
| Methicillin-resistant *S. aureus* | AR0482 | ++ | ++ |
| Methicillin-resistant *S. aureus* | AR0483 | ++ | ++ |
| Methicillin-resistant *S. aureus* | AR0484 | ++ | ++ |
| Methicillin-resistant *S. aureus* | AR0485 | ++ | ++ |
| Methicillin-resistant *S. aureus* | AR0486 | ++ | ++ |
| Methicillin-resistant *S. aureus* | AR0487 | + | -/+ |
| Methicillin-resistant *S. aureus* | AR0488 | + | -/+ |
| Methicillin-resistant *S. aureus* | AR0489 | + | - |
| Methicillin-resistant *S. aureus* | AR0490 | ++ | ++ |
| Methicillin-resistant *S. aureus* | AR0491 | -/+ | - |
| Methicillin-resistant *S. aureus* | AR0492 | + | - |

### II. Variation in the demise – resurrection population dynamics presented in Figure 1B

To explore the generality of the PYO^Sa^ – *S. aureus* Newman serial transfer results presented in Figure 1B, we performed the experiments with ten 2 ml and six 10ml cultures. Both had effectively the same initial densities of phage and bacteria. The dynamics of only one of the ten 2 ml samples serially transferred were similar to the demise–resurrection dynamics observed from Figure 1B. While all six 10 ml serial transfer cultures were turbid by the 5^th^ transfer, some did not become turbid until the 4^th^ transfer. We interpret this observation to be consistent with the hypothesis that the resurrection of these phage-exposed bacteria is a stochastic process in *S. aureus* and thus is more likely to occur in the 10ml populations because the total number of bacteria is 5-fold greater than that in 2 ml cultures.

Also consistent with this stochastic hypothesis is that the time before the recovery of the *S. aureus* population due to the ascent of small colony variants, S, is variable. In Figure 1B, all three serial transfer populations started the recovery to high density by the second transfer. This was not the case for a repeat of this experiment. The recovery of one of the three populations (noted in red) started at the end of the first transfer (Figure S1A).

In Figure 1D, where the serial transfer cultures were initiated with *S. aureus* isolated at the end of serial transfer experiments with the PYO^Sa^, the evolved bacteria, the phage were maintained in one of three cultures and were lost in two. In the repeat of this experiment with three independently evolved strains, the phage were lost in all three (Figure S1B).

**Figure S1.**
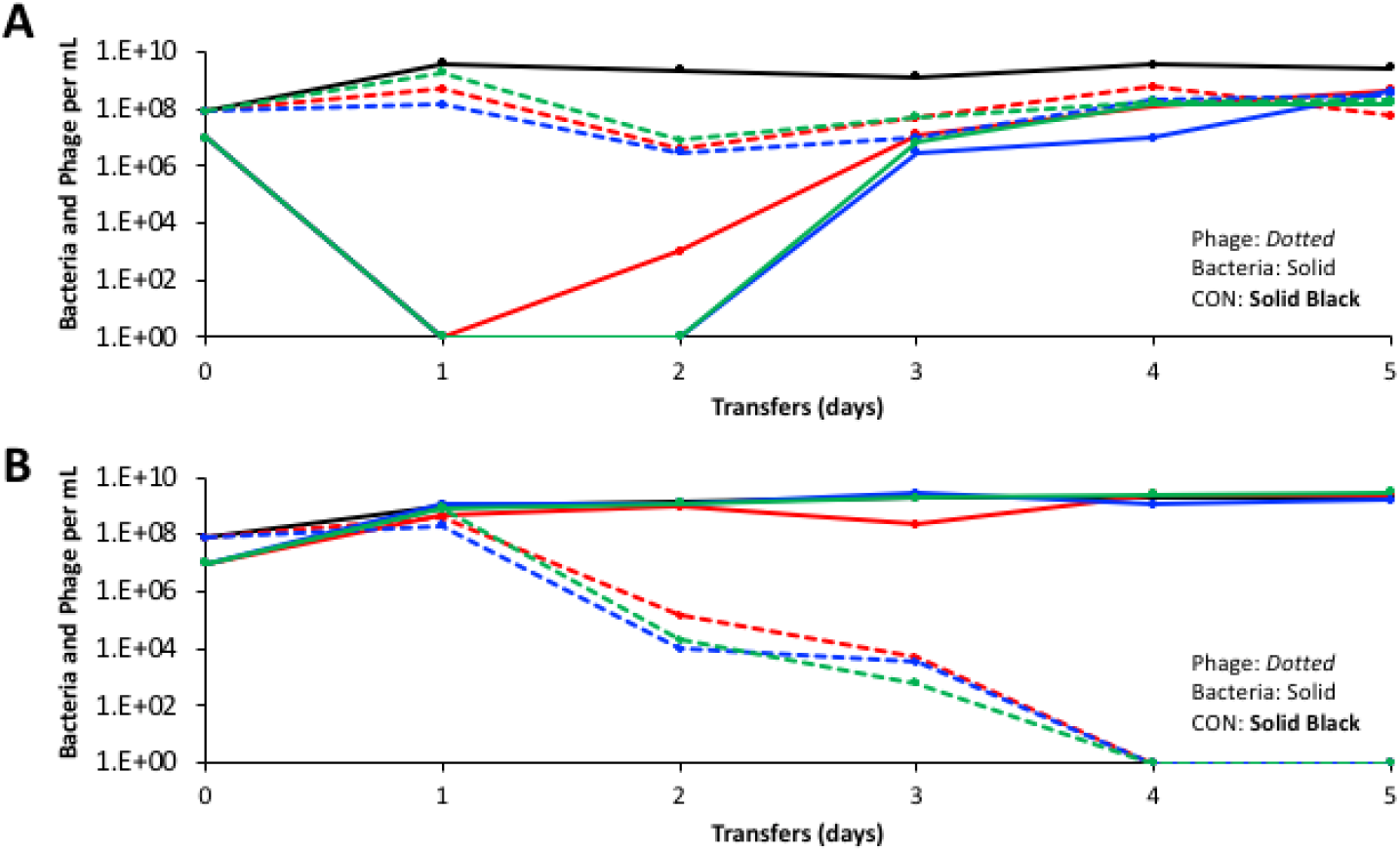
Changes in the densities of bacteria and phage in (1/100) 10 ml serial transfer cultures of *S. aureus* Newman and PYO^Sa^ and a phage-free control, CON. **A)** Three independent cultures initiated with *S. aureus* Newman (W) that had not previously been exposed to PYO^Sa^. **B)** Three independent cultures initiated with *S. aureus* Newman (E) obtained from the 5^th^ transfer of independent serial passage experiments with *S. aureus* Newman mixed with PYO^Sa^.

### III. Genotypes and phenotypes of *S. aureus* mutants selected in the presence of PYO^Sa^

**Table S2.** Small colony variant (S) mutants selected by exposure to PYO^Sa^ in experiments of the type shown in Fig. 1*B*.

| Gene | Mutation |
| --- | --- |
| *femA* | G>T (nt -9) |
| *femA* | G>A (nt -12) |
| *femA* | M1I |
| *femA* | Δ G124 - A126 |
| *femA* | I171AN |
| *femA* | Y327D |
| *femA* | 72 bp Dup (nt 987-1058) |
| *femA* | G331D |
| *femA* | G368V |

Mutations in *femA* were identified after whole genome sequencing of small colony mutants.

**Table S3.**
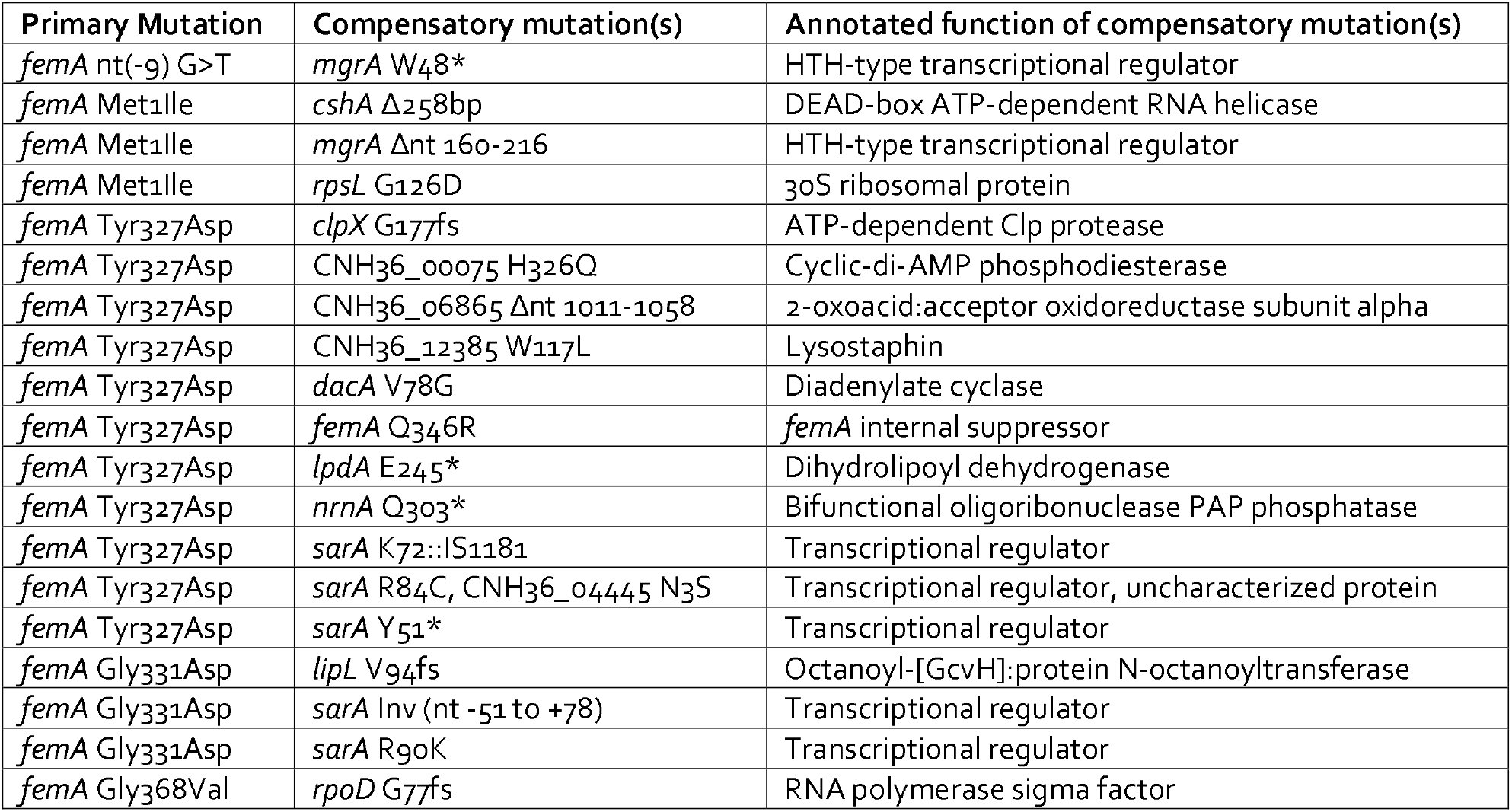
Genotypes of evolved (E) mutants from 5 different small colony variant (S) mutants.

Secondary compensatory mutations associated with the evolved state were identified by whole genome sequencing.

**Table S4.** Relative growth rates and MICs of wild-type (W), small colony variant (S), and evolved (E) compensated mutants.

| Genotype / Mutations |  | Growth Rate | MIC mg/L |  |
| --- | --- | --- | --- | --- |
| Primary | Compensatory | Relative | Lysostaphin | Oxacillin |
| Wild-type | | 1 | $\leq 0.008$ | 2 |
| *femA* M1I |  | 0.53 | 1 | 0.25 |
| *femA* I171N |  | 0.48 | 4 | 0.125 |
| *femA* Y327D |  | 0.59 | >4 | 0.125 |
| *femA* Y327D | *femA* Q346R | 0.76 | 0.125 | 0.063 |
| *femA* Y327D | *sarA* K72::IS1181 | 0.93 | >4 | 0.125 |
| *femA* G331D |  | 0.53 | >4 | 0.063 |
| *femA* G331D | *sarA* INV (nt -51-+78) | 0.94 | >4 | 0.125 |

Doubling time for wild-type is 25.8 min.

### IV. A hypothesis to account for the population and evolution dynamics of the *S. aureus* and PYO^Sa^

In our article we postulated that the dynamics presented in Figure 1B can be attributed to the generation of small colonies from the wild-type *S. aureus* Newman and their ascent due to selection mediated by PYO^Sa^ phage. While the PYO^Sa^ phage adsorb to these small colony variants, they do not kill them, and the infecting phage are lost. The phage are maintained because the small colony variants are of low fitness and continually produce normal or near normal colony variants, an “evolved” state, that can support the replication of the phage. These evolved cells are not stable and continually produce small colonies. This hypothesis is illustrated in Figure S2.

**Figure S2.**
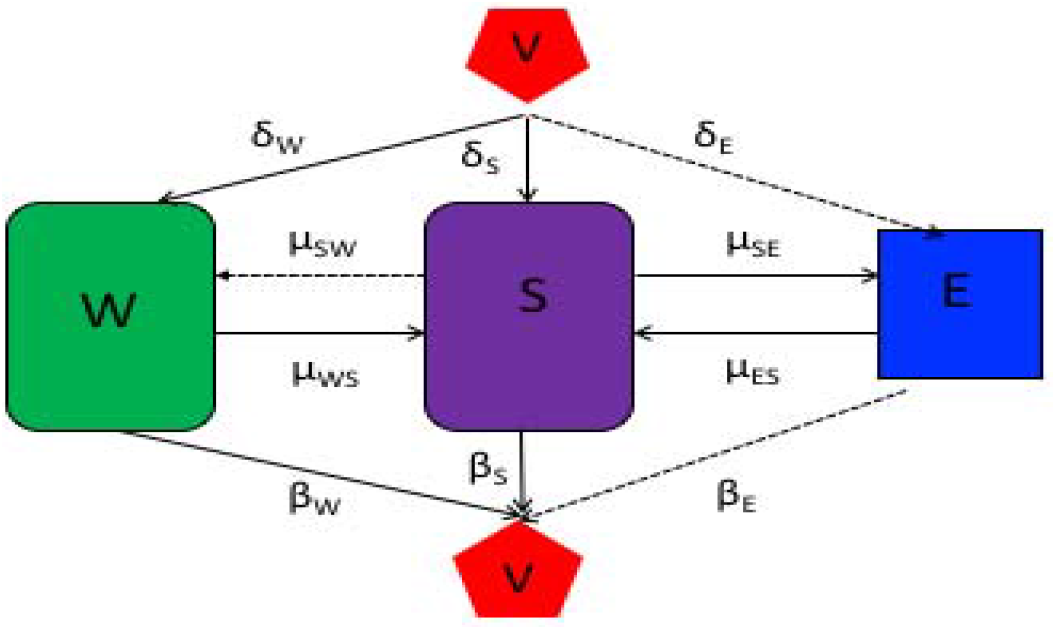
A population dynamic model to account for the observed changes in the densities of bacteria and phage in Figure 1B. The variables, W, S and E are, respectively the wildtype *S. aureus* Newman, small colonies, and the evolved bacteria, cells per ml, and V the PYO^Sa^ phage, particles per ml. The parameters δ_W_, δ_S_ and δ_E_ are the adsorption rate constants per cell per ml. The parameters β_W_, β_S_ and βE are the number of phage particles produced per infected cell burst. We assume that β_S_ =0, the S are infected by PYO^Sa^ but the phage but do not replicate or kill these bacteria. The parameters μ_SW_, μ_WS_, μ_SE_ and μ_ES_ are the transition rates, per cell per hour, between the different states.

The population is in liquid culture in which there is a limiting resource of concentration, r, μg/ml. The net growth rate of the bacteria is proportional to its maximum rate of growth and a hyperbolic function of concentration of the limiting resource, r, μg/ml and a constant, k_i_ which is the concentration of the resource when the growth rate is half its maximum value., where the subscript i is W, S or E (1). We assume there is no latent period and upon adsorption, W and E are killed by the phage and instantly produce, β_W_ and βE phage particles per cell (2). Although the phage adsorb to the small colonies, they are not killed. As the bacteria grow the concentration of the resource declines at a rate proportional to the net growth rates of the bacteria, and a conversion efficiency parameter, *e_i_* μg/ml, the conversion efficiency, which is the amount of resource needed to produce a new cell of that type (3). To account for the declining physiological state of the bacteria as the bacteria approach stationary phase, r=0, we assume that the rates of phage adsorption and transitions between states is proportional to the net growth rate of that cell line. With these definitions and assumptions, the rates of change in the densities of the bacteria of different states, the density of free phage, and the concentration of the resource are given by the below set of coupled differential equations.

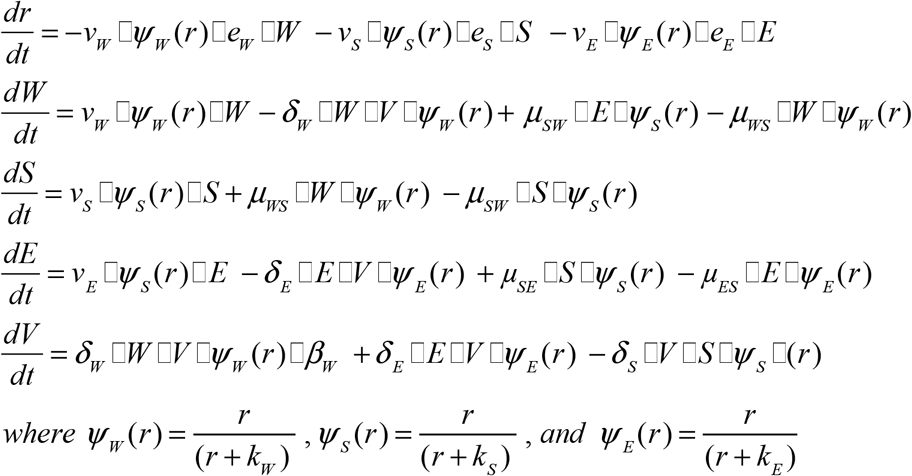

#### Numerical Solutions – Simulations

To solve this set of coupled differential equations and those for the models that follow we use Berkeley Madonna. The population growth and phage infection parameters employed for these numerical solutions, simulations, are of the range estimated for PYO^Sa^ and *S. aureus* Newman in MHII medium. To simulate a serial transfer mode of population maintenance every 24 hours there is a 100-fold reduction in densities of the bacteria and phage and the resource concentration is restored to its maximum level of 1000μg per ml. For copies of the program and instructions for it use write to.

#### Simulation Results

In Figure S3, we follow the changes in the densities of the bacteria and phage in simulated serial populations. As observed in Figure 1B, the model predicts, as seen in Figure S3A, that upon the first encounter with the phage, V, the density of susceptible cells, W, and the total cell density of the bacteria, NT, will declines whilst that of the phage increase. In subsequent transfers, the population of bacteria recovers and becomes dominated by small colonies. As a consequence of the transitions between the different states of bacteria, the phage and all three bacterial populations are maintained with small colonies dominating the bacterial community. If we make the small colonies less efficient in the use of the resource, e_c_ =5×10^−6^ rather than 5×10^−7^, the total density of the small colony population is lower (Figure S3A). When cultures of just small colonies are started without the phage, they are ultimately diluted out due to their lower growth rate and low production rate in the absence of the phage and are overtaken by the evolved population being generated through reversion from small colonies (Figure S3B). When the evolved cells and phage are mixed, as observed in Figure 1D, the population recovers to full density more rapidly than when sensitive cells and phage are mixed (Figure S3C). However, in these simulations, the phage continue to be maintained, which was the case for only one of the two parallel experiments in the main body of the paper and in none of the parallel experiments presented in supplemental Figure S1B. This model can also account for why the mixture of the ancestral *S. aureus* Newman and small colony variants can maintain the phage (Figure 1E) with little effect on the density of bacteria (Figure S3D).

**Figure S3.**
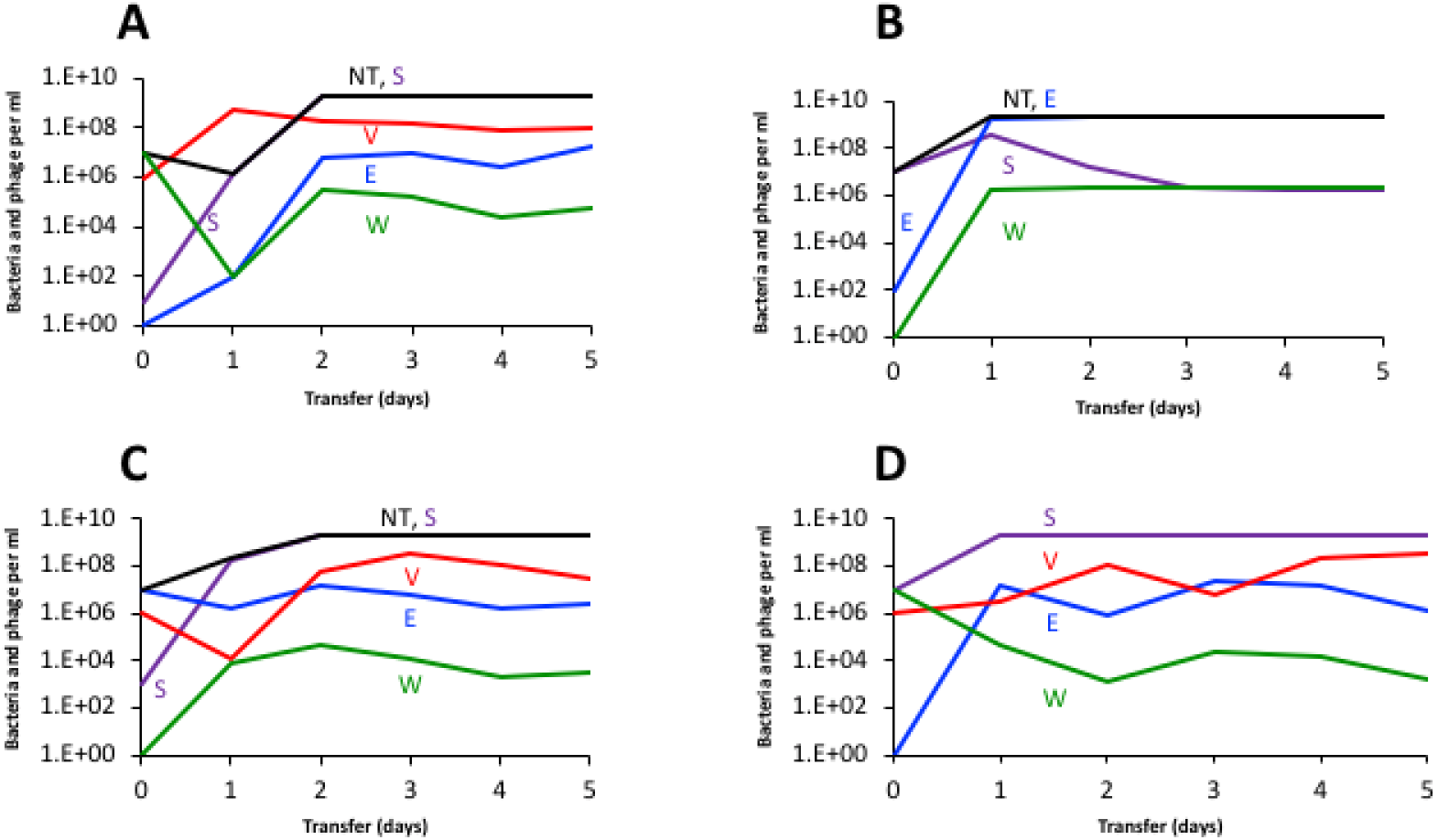
Simulated serial transfer populations. Changes in the densities of bacteria and phage in serial transfer culture. Standard parameters, v_W_= v_E_=1.7, v_S_=0.5, δ_W_= δ_E_=δ_S_= 2×10^−7^, β_W_-β_E_= 80, β_S_=0, e_w_ = e_e_=5×10^−7^, ec=5E×10^−6^, C=1000, k_W_ =k_E_=1, k_S_ =10, μ_WS_=10^−6^, μ_SW_=10^−6^, μ_ES_ = 10^−3^, μ_SE_ =10^−3^. NT is the total density of bacteria. **A)** Serial transfer population with W and V and few S and E. **B)** Serial transfer population initiated with S and miniority populations of W and E, but no phage. **C)** Serial transfer population initiated with E and phage and minority populations of S and W. **D)** Serial transfer populations initiated with a mixture of W, S, and V and a minority of E

### V The joint action of bactericidal antibiotics and phage

In our article, we postulate that the reason bactericidal antibiotics in combination with phage do worse than phage alone (Figure 4) can be attributed to the antibiotics reducing the density of bacteria and thereby the capacity of the phage to replicate. We illustrate this with the following model of the joint action of antibiotics and a bactericidal antibiotic.

#### Model of the joint action of antibiotics and phage

There are two populations of bacteria enumerated in cells per ml. The populations are respectively W (wild-type, which are sensitive to antibiotics) and P (persisters which are sensitive to the phage but phenotypically resistant to antibiotics). There is a single antibiotic of concentration A in μg/ml, a lytic phage of density V particles per ml, and a limiting resource r in μg/ml. The sensitive bacteria grow at a maximum rate, v_N_, with the net growth/death rate ψ(A,r) being proportional to the concentration of the antibiotic and the limiting resource voila,

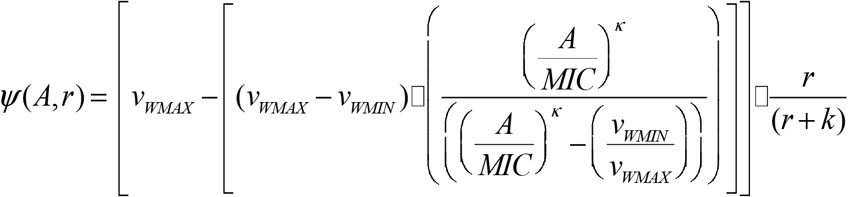

Where v_WMAX_ (>0) is the maximum growth rate, v_WMIN_ (<0) is the minimum growth rate/maximum kill rate, MIC the minimum inhibitory concentration of the antibiotic, and a shape parameter, κ, such that the greater the value of κ the more acute the function.

The persisters, P, are non-replicating bacteria that are resistant to the antibiotic. The phage adsorb to the persisters but do not replicate on them. The rate constant of adsorption of the phage to sensitive cells is δ_W_ and that to the persisters δ_P_. Infections of phage to sensitive cells produce β phage particles per cell. With a rate y per cell per hour, sensitive cells produce persisters, W-->P, and with a rate x per cell per hour, persisters produce sensitive cells, P -->W. With these definitions and assumptions, the rates of change in the densities of phage and the concentration of the resources are given by

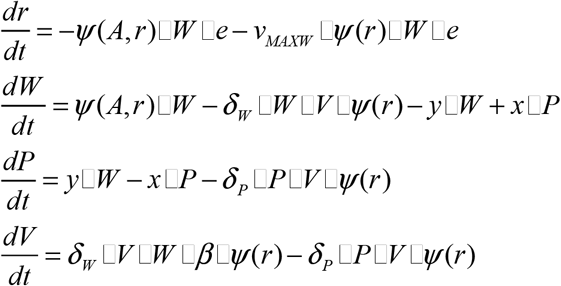

*where* 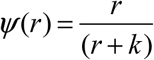 *and e μg is the amount of the* lim*iting resource need to produce a new cell*

#### Numerical Solutions – Simulations

To solve these equations and simulate the dynamics, we use Berkeley Madonna. Copies of the programs are available from.

##### Results

In Figure S4, we illustrate the rate of change in the total density of bacteria with a bactericidal antibiotic alone, with phage alone, and with the phage and antibiotic together. The rate of decline in the density of bacteria is lowest in the simulations where antibiotics are used alone. The highest rate of decline in the density of bacteria is obtained in the simulation where phage are used alone. When phage are used in combination with bactericidal antibiotics, they do not increase in density to the same extent that they do in the absence of the antibiotics. The leveling off in the density of bacteria is a consequence of persistence (4–6).

**Figure S4.**
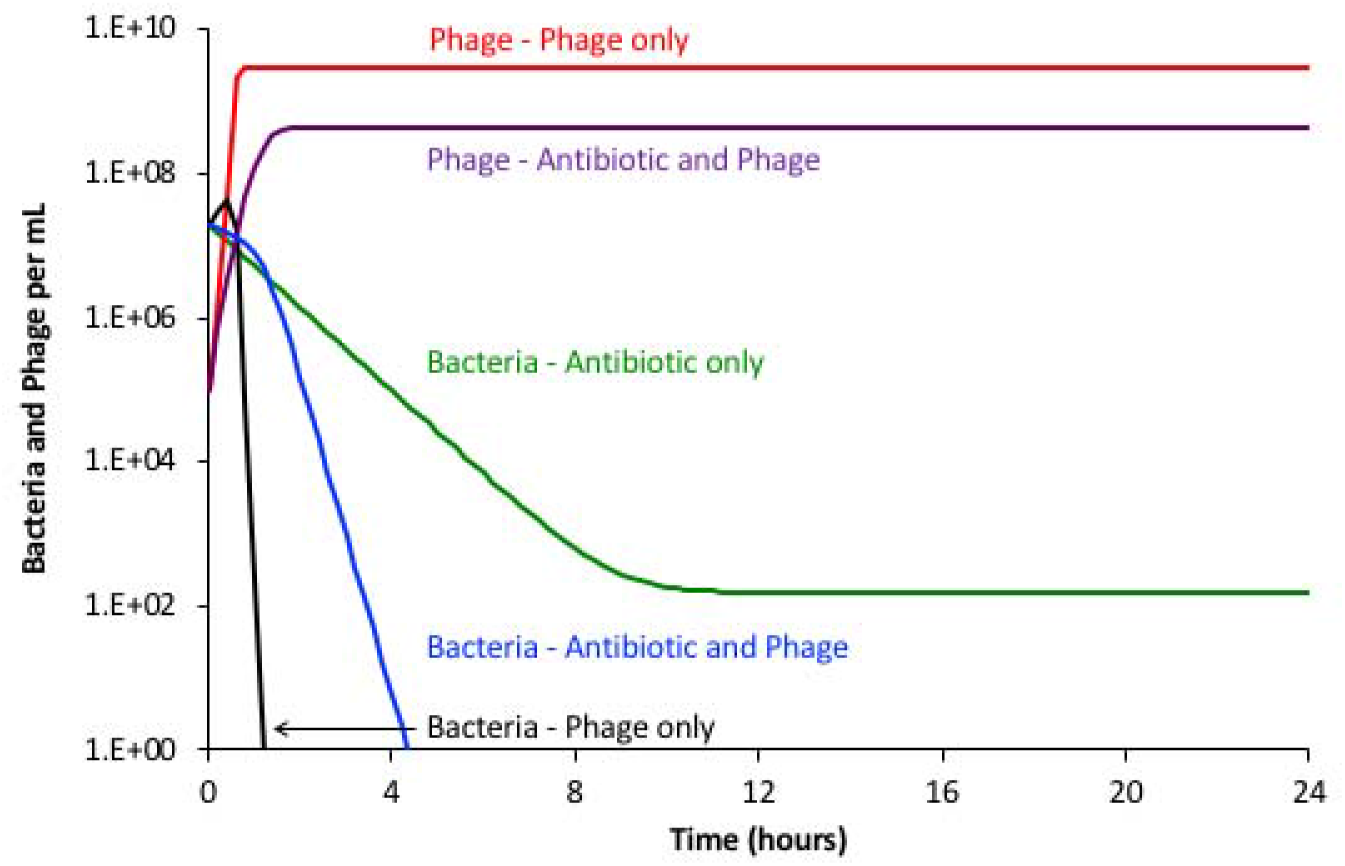
Simulation results: changes in the densities of bacteria and phage with different treatments. Parameter values. v_SMAX_=2.0, v_SMIN_ −2.0, κ=1.0, MIC=1.0, e=5×10^−7^, k=1.0, x=y=10^−5^, δ_P_=10^−8^, δ_S_=10^−8^, β=50, A =2 μg/ml, r(0)=1000.

### VI. Persistence and heteroresistance

In the absence of phage in serial transfer culture, the super MIC concentration of oxacillin does not eliminate the bacteria but rather maintains the density at levels markedly lower than the antibiotic-free controls (Figure 4 A center). We postulate that this can be attributed to persistence in serial transfer culture. In the absence of phage, ciprofloxacin initially reduces the density of *S. aureus* but in subsequent transfers, the density of these bacteria return to levels within the range anticipated for antibiotic-free controls (Figure 4A bottom). We postulated that these pharmacodynamics can be attributed to heteroresistance (7). In the following, we present the evidence for persistence for oxacillin-treated cultures and heteroresistance for ciprofloxacin-treated cultures. Using a mathematical - computer simulation model, we illustrate how these postulated mechanisms can account for the observed pharmacodynamics of these drugs with *S. aureus* Newman.

#### A model for antibiotics in serial transfer culture with persistence and heteroresistance

There are three populations of bacteria, wild-type, heteroresistant, and persisters with designations and densities, W, H, and P cells per ml; a single antibiotic of concentration, A μg/ml; and a limiting resource of concentration r μg/ml. As in our model of the joint action of antibiotics and phage (II), we assume that the rate of growth, of bacteria of each of these states, ψ_i_(A,r) is a product of a Hill function for the antibiotic (4), and a Monod function for the resource (1).

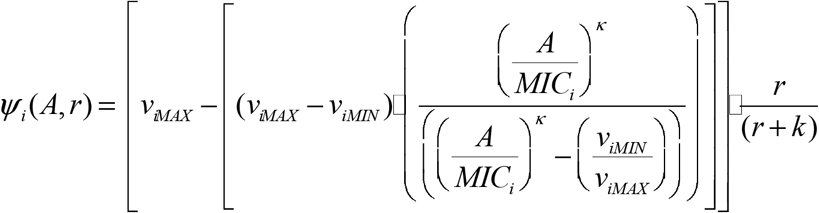

The subscript i represents the bacterial population, W, H, or P. For strain i, v_iMAX_ (>0) is the maximum growth rate, v_i*MIN*_ (<0) the minimum growth rate/maximum kill rate, and MIC_i_ the minimum inhibitory concentration of the antibiotic. Bacteria of the same state have the same Hill coefficient and Monod constant, respectively κ and k.

In these simulations, we assume the persisters can replicate, v_PMAX_>0, and are resistant to the antibiotics, v_PMIN_=0. As in (7), the MIC for the antibiotic of the heteroresistant cells is greater than the sensitive, MIC_H_ >MIC_S_. There is a transition from W to P and from P to W respectively at rates, x and y per cell per hour, and a transition from W to H and H to W at rates, x_H_, and y_H_ per cell per hour. Finally, we assume that the concentration of the antibiotic can decline at a rate d per hour. With these definitions and assumptions, the rates of change in the densities of the bacterial population and concentration of the antibiotic and resource are given by:

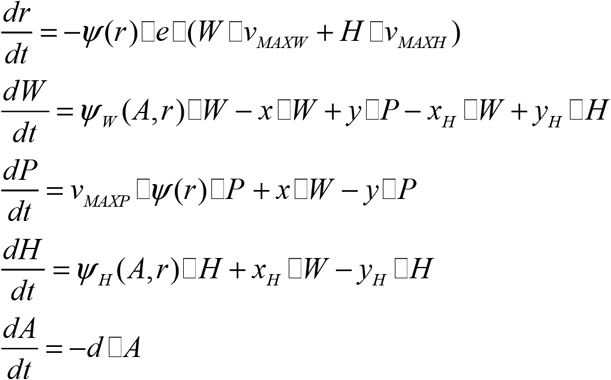

*where* 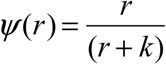

#### Numerical Solutions

To solve these equations, we simulate the changes in the densities of the bacterial populations and changes in the concentration of the antibiotics with Berkeley Madonna. As in our experiments, in these simulations every 24 hours the density of the bacteria is reduced by a factor of 100, and new resources and antibiotics are added.

**Figure S5.**
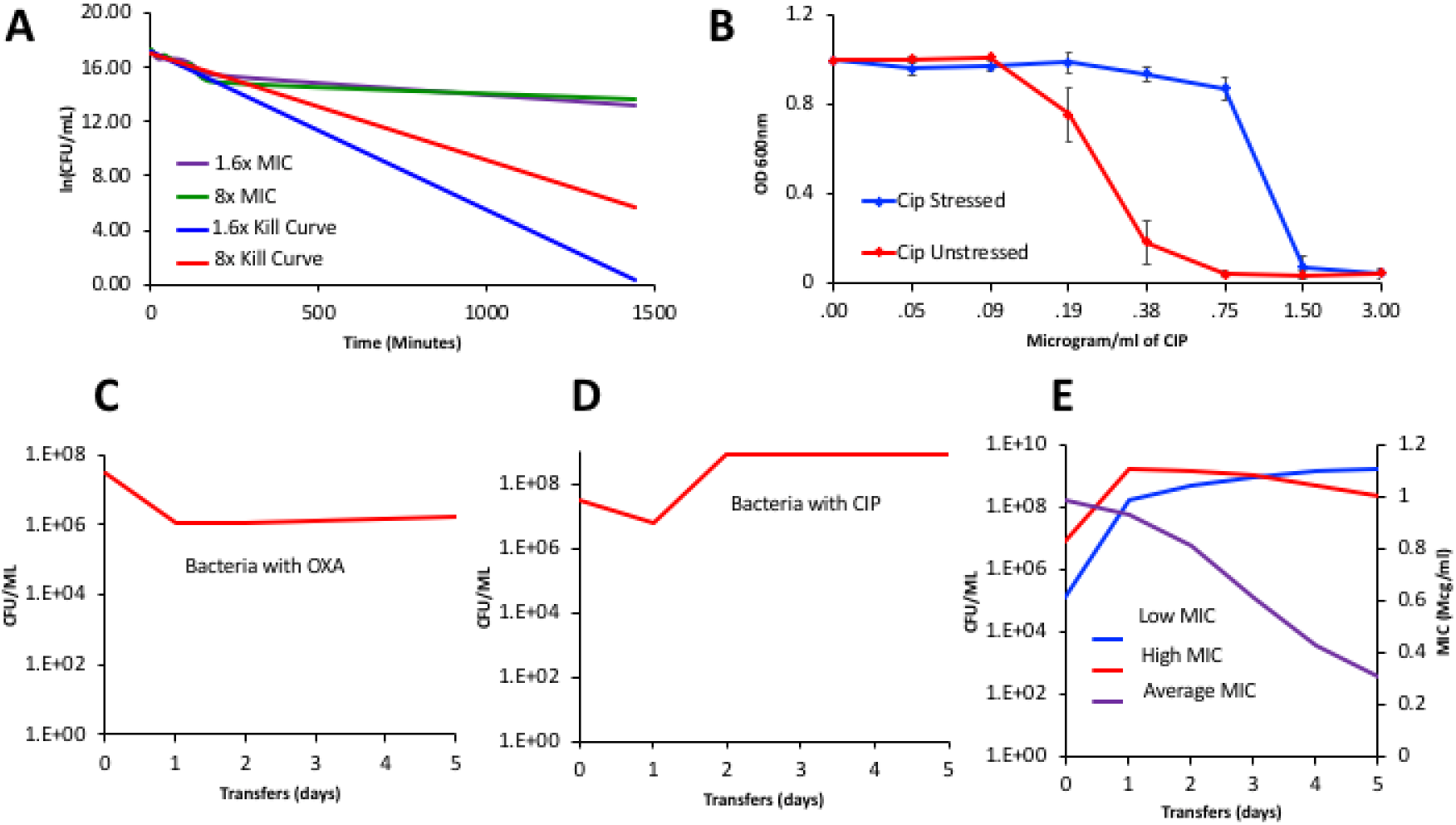
*S. aureus* antibiotic pharmacodynamics **A)** Natural log of the change in density of growing cultures of *S. aureus* Newman exposed to 1.6X and 8X MIC, low and high for 24 hours (1440 minutes). The green and purple lines are experimental changes in density estimated by plating; while the blue and red lines are the results of a linear regression of the changes in density estimated during the first 180 minutes of exposure. **B)** Changes in the optical density of experimental cultures of *S. aureus* Newman exposed to 5μg/ml ciprofloxacin, mean and standard error of the ODs for three replicas. In red are cells that were treated with ciprofloxacin and then transferred without the presence of this drug for seven days before the MIC determination was performed. In blue are cells that were treated with ciprofloxacin for one week before the MIC was performed. **C, D, E)** Simulation Results Common parameters: v_MAXS_=1.5, v_MINS_=-0.6,v_MAXSR_ = 1.2, v_MINSR_=-3, ks=1,kh=1,k=1,e=5.0E-7 **C)** Changes in the densities of bacteria in serial transfer culture with persistence with parameters in the range estimated for *S. aureus* in oxacillin. Specific parameters: vp=0.2, MIC_S_= 1.8, x_P_=1.0e-4, y_P_=1E-3, A_MAX_=3. Exposure to oxacillin does not start until the second transfer (compare to Figure 4A center). **D) and E)** Heteroresistance for ciprofloxacin MIC_S_= 0.218, MIC_H_=1, x= 1E-6, y=1e-3, x_P_=0, y_P_=0, A_MAX_=0.5 Changes in the densities of bacteria in serial transfer culture with heteroresistance with parameters in the range estimated for *S. aureus* in ciprofloxacin. Exposure to ciprofloxacin doesn’t start until the second transfer (compare to Figure 4A bottom). **E)** Unique Parameters for E: MIC_S_= 0.218, MIC_H_=1, x=1E-6, 1E-3 x_P_=0, y_P_=0. Simulation of the changes in the densities of bacteria and average MIC of the antibiotic mixture of the low MIC sensitive and the high MIC heteroresistant populations in serial transfer culture in the absence of the antibiotics.

In Figure S5A, we present the results of time-kill experiments with *S. aureus* Newman exposed to oxacillin. The observation that the rate of decline in the viable density of sensitive cells during the first few hours of exposure considerably exceeds the rate of decline later in the experiment is what would be anticipated for persistence. In Figure S5B, we present the evidence for heteroresistance; higher concentrations of ciprofloxacin are needed to kill cultures initiated with bacteria exposed to ciprofloxacin than to kill bacteria cultured in the absence of this drug. If we allow for persistence with parameters in the range estimated for oxacillin the predicted changes in the density of *S. aureus* Newman in serial transfer culture with oxacillin are similar to those observed (compare Figures S4A and S4C). If we allow for heteroresistance with the parameter in the range estimated for *S. aureus* Newman in serial transfer culture with ciprofloxacin the results are similar to those observed (compare figures S5B and S5D). One property of heteroresistance is that when the bacteria are removed from the drug, the resistant population will decline in frequency, and the average MIC will decline to levels similar to that of the original sensitive population (7), see Figure S5E.

